# The lemur baseline: How lemurs compare to monkeys and apes in the Primate Cognition Test Battery

**DOI:** 10.1101/2020.04.21.052852

**Authors:** Claudia Fichtel, Klara Dinter, Peter M. Kappeler

**Affiliations:** Behavioral Ecology & Sociobiology Unit, German Primate Center (DPZ), Kellnerweg 4, 37077 Göttingen, Germany; Leibniz ScienceCampus “Primate Cognition”, Göttingen, Germany; Department of Sociobiology/Anthropology, Johann-Friedrich-Blumenbach Institute of Zoology and Anthropology, University of Göttingen, Kellnerweg 6, 37077 Göttingen, Germany

**Keywords:** cognition, Primate Cognition Test Battery, primates, lemurs

## Abstract

Primates have relatively larger brains than other mammals even though brain tissue is energetically costly. Comparative studies of variation in cognitive skills allow testing of evolutionary hypotheses addressing socioecological factors driving the evolution of primate brain size. However, data on cognitive abilities for meaningful interspecific comparisons are only available for haplorhine primates (great apes, Old- and New World monkeys) although strepsirrhine primates (lemurs and lorises) serve as the best living models of ancestral primate cognitive skills, linking primates to other mammals. To begin filling this gap, we tested members of three lemur species (*Microcebus murinus, Varecia variegata, Lemur catta*) with the Primate Cognition Test Battery, a comprehensive set of experiments addressing physical and social cognitive skills that has previously been used in studies of haplorhines. We found no significant differences in cognitive performance among lemur species and, surprisingly, their average performance was not different from that of haplorhines in many aspects. Specifically, lemurs’ overall performance was inferior in the physical domain but matched that of haplorhines in the social domain. These results question a clear-cut link between brain size and cognitive skills, suggesting a more domain-specific distribution of cognitive abilities in primates, and indicate more continuity in cognitive abilities across primate lineages than previously thought.

## INTRODUCTION

One central question in comparative cognition is why primates have evolved larger brains and enhanced cognitive skills compared to other equally-sized mammalian species (Shettleworth 2010). Among primates, this effect is paralleled by a disproportionate increase in brain size from strepsirrhines to haplorhines and humans (Dunbar 1992; Isler et al. 2008; Jerison 1973; Martin 1981). Because larger brains are energetically more expensive (Aiello and Wheeler 1995), they are assumed to confer benefits with regard to enhanced cognitive abilities that compensate this additional investment (Navarrete et al. 2011; Reader and Laland 2002; Reader et al. 2011).

Several non-mutually exclusive hypotheses on the evolution of brain size have been proposed to account for the distinctive cognitive abilities of primates (Dunbar and Shultz 2017). According to the *General intelligence hypothesis*, larger brains are thought to confer an advantage because of faster learning and larger memory capacities (Spearman 1904). The *Ecological intelligence hypothesis* suggests that environmental and ecological challenges in food acquisition, including spatial and spatio-temporal processes to memorize seasonally available food or manipulative skills for extractive foraging, selected for larger brains (Byrne 1996; Clutton-Brock and Harvey 1980; Heldstab et al. 2016; Milton 1981; Powell et al. 2017). Several versions of the *Social brain hypothesis* posit that increased cognitive skills in primates evolved in response to the constant challenges associated with the complexity of social life, such as competition and cooperation within larger social groups (Byrne and Whiten 1988; Dunbar 1992; Dunbar and Shultz 2007; Humphrey 1976; Jolly 1966a; Kudo and Dunbar 2001). However, support for the *Social brain hypothesis* is not uniform in other taxa, with brain size correlating positively with measures of sociality in some insectivores, bats and ungulates (e.g. Barton et al. 1995; Byrne and Bates 2010; Dunbar and Bever 1998; Shultz and Dunbar 2006), but not in corvids (Emery et al. 2007; Shultz and Dunbar 2007), and it is equivocal in carnivores (Benson-Amram et al. 2016; Dunbar and Bever 1998; Finarelli and Flynn 2009; Holekamp et al. 2007; Pérez-Barbería et al. 2007). Moreover, recent comparative analyses among primates indicated that brain size is associated with ecological (home range size, diet, activity period), but not with social factors (DeCasien et al. 2017; Powell et al. 2017), also challenging the social brain hypothesis.

Since these studies usually link interspecific variation in brain size with certain socio-ecological factors, it is essential to understand how brain size actually impacts cognitive skills. Hence, comparative studies of cognitive abilities, ideally using identical tests, across the primate order and beyond are required. However, comparisons of performance in cognitive experiments across species may fail due to variation in the experimental set-up and specific methods (van Horik and Emery 2011; Krasheninnikova et al. 2019; MacLean et al. 2012).

To overcome this problem, Herrmann and colleagues (2007) assembled a systematic toolbox for comparative analysis, called the *Primate Cognition Test Battery* (PCTB), which compared cognitive skills in various tasks in the physical and social domain among 2.5-year-old children, chimpanzees (*Pan troglodytes*) and orangutans (*Pongo pygmaeus*). The physical domain deals with the spatial-temporal-causal relations of inanimate objects, while the social domain deals with the intentional actions, perceptions, and knowledge of other animate beings (Tomasello and Call 1997). These tests revealed that children and chimpanzees have similar cognitive skills for dealing with the physical world, but children have increased cognitive skills for dealing with the social world, particularly in the scale of social learning. These results support the *Cultural intelligence hypothesis*, a variant of the Social brain hypothesis, suggesting that exchanging knowledge within human cultural groups requires specific socio-cognitive skills, such as social learning or Theory of Mind (e.g. Boyd and Richerson 1998; Herrmann et al. 2007; Whiten and van Schaik 2007).

Application of the PCTB to two other haplorhine primate species, long-tailed macaques (*Macaca fascicularis*) and olive baboons (*Papio anubis*), revealed that both species performed similarly to great apes in both the physical and the social domain (Schmitt et al. 2012). Specifically, chimpanzees outperformed macaques only in tasks on spatial understanding and tool use. Since chimpanzees have relatively larger brains than macaques or baboons (Isler et al. 2008; Jerison 1973), these results question the clear-cut relationship between cognitive performance and brain size (Schmitt et al. 2012). In addition, four closely related macaque species that differ in their degree of social tolerance, performed similarly in cognitive tests of the PCTB in the physical domain. However, socially more tolerant species performed better in one task of the social domain and the inhibitory control task, suggesting that social tolerance is associated with a set of cognitive skills that are specifically required for cooperation (Joly et al. 2017). Thus, further studies on additional non-human primates are required to explore the interrelationships among cognitive abilities, socio-ecological traits and brain size (ManyPrimates et al. 2019).

Strepsirrhine primates are the obvious candidates for such an extended comparative approach because they represent the best living models of the earliest primates and the link between primates and other mammalian orders (Fichtel and Kappeler 2010; MacLean et al. 2008). Strepsirrhines split off from the main primate lineage approximately 60 million years ago and retained many ancestral primate traits (Martin 1990; Yoder et al. 1996; Yoder and Yang 2004). Importantly, strepsirrhine primates have relatively smaller brains than haplorhines, and their brain size does not correlate with group size (MacLean et al. 2009). Although older studies suggested that strepsirrhine primates possess physical cognitive abilities that are inferior to those of haplorhines (e.g. Ehrlich et al. 1976; Jolly 1964; Maslow and Harlow 1932), recent studies indicated that their cognitive skills are similar to those of haplorhines (e.g. Deppe et al. 2009; Fichtel and Kappeler 2010; Kittler et al. 2015, 2018; Santos et al. 2005a, b). However, existing studies of strepsirrhine cognition used isolated tests, hampering systematic interspecific comparisons. Hence, a comprehensive study investigating a broad variety of tasks addressing different cognitive skills in lemurs, and replicating the exact same methods used in the PCTB, seems indicated for a systematic comparison across both primate suborders.

To this end, we applied the PCTB to three species of lemur that differ in key socio-ecological traits: ring-tailed lemurs (*Lemur catta*), black-and-white ruffed lemurs (*Varecia variegata*; in the following: ruffed lemurs) and gray mouse lemurs (*Microcebus murinus*, Table 1). Mouse lemurs have one of the smallest brain sizes among primates, and absolute brain size increases from mouse lemurs over ring-tailed lemurs to ruffed lemurs (Isler et al. 2008). Ring-tailed lemurs are diurnal opportunistic omnivores that live in groups of on average 14 individuals (Gould et al. 2003; Jolly 1966b; Sussman 1991). Ruffed lemurs are diurnal, frugivorous and live in small groups (average 6 individuals), exhibiting a fission-fusion social organization (Baden et al. 2015; Holmes et al. 2016; Vasey 2003). Gray mouse lemurs are nocturnal, omnivorous solitary foragers that form sleeping-groups among related females (Eberle and Kappeler 2006; Isler et al. 2008).

**Table 1.** Summary of the most important traits for the seven non-human primate species.

| species | n | ECV<br>(cc) | %<br>fruit | dietary<br>breadth | social<br>system | average<br>group size |
| --- | --- | --- | --- | --- | --- | --- |
| **chimpanzees**<br>(*Pan troglodytes*) | 106 | 368.4 | 66 | 6 | group | 47.6 |
| **orangutans**<br>(*Pongo pygmaeus*) | 32 | 377.4 | 64 | 6 | solitary | 1.5 |
| **olive baboons**<br>(*Papio anubis*) | 5 | 167.4 | 62 | 6 | group | 69 |
| **long-tailed**<br>**macaques**<br>(*Macaca fascicularis*) | 10-13 | 64 | 66.9 | 5 | group | 26 |
| **ruffed lemurs**<br>(*Varecia variegata*) | 13 | 32.1 | 92 | 4 | group | 6 |
| **ring-tailed lemurs**<br>(*Lemur catta*) | 26-27 | 22.9 | 54 | 5 | group | 11 |
| **grey mouse lemurs**<br>(*Microcebus murinus*) | 9-16 | 1.6 | 31.3 | 4 | solitary | 1 |

n=number of individuals, ECV=endocranial volume (absolute brain size), % fruit=percentage of fruit in the diet; Data from: Herrmann et al. 2007; Schmitt et al. 2012; Isler et al. 2008; MacLean et al. 2014; Dammhan and Kappeler 2006; Radespiel et al. 2006; Lahann 2007.

According to the *General intelligence hypothesis*, we predicted that the tested apes and monkeys outperform lemurs because they have absolutely larger brains (Table 1). In accordance with the *Ecological intelligence hypothesis* we predicted that the more frugivorous species or those with a broader dietary breadth perform better (Table 1). Because lemurs generally live in smaller groups than monkeys and apes (Kappeler and Heymann 1996), we predicted that they should have inferior cognitive abilities than the already tested group-living species according to the *Social intelligence hypothesis* (Table 1).

## METHODS

Experiments were conducted with adult individuals of gray mouse lemurs (n=9-15), ring-tailed lemurs (n=26-27) and black-and-white ruffed lemurs (n=13). All individuals were born in captivity and housed in enriched or semi-natural environments, either at the German Primate Centre (DPZ, Göttingen) or the Affenwald Wildlife Park (Straußberg, Germany). The lemurs at the Affenwald range freely within a 3.5 ha natural forest enclosure. At the DPZ, ring-tailed and ruffed lemurs are offered indoor and outdoor enclosures equipped with enriching climbing materials and natural vegetation. The nocturnal mouse lemurs are kept indoors with an artificially reversed day-night-cycle, and cages are equipped with climbing material, fresh natural branches and leaves. All individuals were tested individually in their familiar indoor enclosures and were unfamiliar with the presented tasks. Since some individuals passed away during the course of the study, not all individuals participated in every task of the test battery (Table S1, Supplemental). To ensure comparability with the previous studies, the experimental setup was replicated after the PCTB (Herrmann et al. 2007; Schmitt et al. 2012), and only objects presented in the tests were adjusted in size for lemurs.

### Ethical statement

All animal work followed relevant national and international guidelines. The animals were kept under conditions documented in the European Directive 2010/63/EU (directive on the protection of animals used for experimental and other scientific purposes) and the EU Recommendations 2007/526/EG (guidelines for the accommodations and care of animals used for experimental and other scientific purposes). Consultation and approval of the experimental protocols by the Animal Welfare Body of the German Primate Center is documented (E2-17).

### General testing procedure

During the experiments, individuals were briefly separated from the group. The testing apparatus for all tasks consisted of a table with a sliding board on top that was attached to the fence of the subjects’ enclosures (Figure S2, Supplemental). In most of the tasks two or three opaque cups (ruffed- & ring-tailed lemurs: Ø 6.8 cm x 7.5 cm; mouse lemurs: Ø 2.5 cm x 3 cm), which were placed upside down in a row on the sliding board, were used to cover the food reward (see also Supplemental). If necessary, a cardboard occluder was put on top of the sliding board between the experimental setup and the individual to hide the baiting process from the individuals. The position of the reward was randomized and counter-balanced across all possible locations, and the reward was never put in the same place for more than two consecutive trials. Once the board was pushed into reach of an individual, the experiment began and, depending on the task, the individual had to manipulate an item or indicate its choice by pointing or reaching towards the chosen item, to obtain the reward if chosen correctly. If the choice was incorrect, the correct location of the reward was shown to the individual after each trial.

For most of the tasks at least 6 trials were conducted per individual and setup (Table S1, Supplemental). Raisins and pieces of banana served as rewards. During testing, no possible cues to where the reward was located were provided by the experimenter; she simply put her hands on her lap and her gaze was directed downwards. All experiments were videotaped and responses of the subjects to the tasks coded afterwards from the videos. A naïve second observer additionally scored 20% of all trials a second time to assess inter-observer reliability. The Interclass Correlation Coefficient was excellent (ICC = 0.985).

### The Primate Cognition Test Battery

All experimental setups and methods were replicated from the *PCTB* (Herrmann et al. 2007; Schmitt et al. 2012). Following Schmitt et al. (2012), we also doubled the number of trials for all object-choice tasks of the test battery (Table S1, Supplemental) to evenly distribute objects between all possible spatial positions and combinations of manipulations. In total, the PCTB consists of 16 different experimental tasks, 10 investigating physical and 6 social cognitive skills. These tasks can be grouped into 6 different scales: space, quantity and causality for the physical and social learning, communication and Theory of Mind for the social domain.

In the *physical domain*, the *scale space* examines the ability to track objects in space in four tasks: spatial memory, object permanence, rotation and transposition. The *scale quantity* tests the numerical understanding of individuals and consists of two tasks: relative numbers and addition numbers. The *scale causality* consists of four tasks: noise, shape, tool use and tool properties to examine the ability to understand spatial-causal relationships. In the *social domain*, the *scale social learning* examines in one task whether individuals use social information provided by a human demonstrator to solve a problem. The *scale communication* examines whether individuals are able to understand communicative cues given by humans in three tasks: comprehension, pointing cups and attentional state. Finally, in the *scale Theory of Mind*, individuals were confronted with two tasks: gaze following and intentions. A detailed description of the general setup and the methodology of the experiments can be found in the supplementary material (Supplemental).

### Temperament, inhibitory control, rank and learning effect

To assess the influence of temperament, inhibitory control and dominance rank on lemurs’ performances in the test battery, individuals participated in a set of additional tests (Herrmann et al. 2007; Schmitt et al. 2012). Due to logistic constraints, the temperament tests could only be conducted with ring-tailed and ruffed lemurs. For temperament, we measured whether individuals would approach novel objects, people and foods (for details see Supplemental). Inhibitory control was measured during an additional session of the spatial memory task, in which out of three cups only the two outer ones were baited with a reward and hence, individuals had to skip the cup in the middle. Dominance rank (high, middle or low-ranking) was inferred by focal observations of ring-tailed and ruffed lemurs but not for the solitary mouse lemurs, according to Pereira and Kappeler (1997). We also controlled for potential learning effects within the trials of a task by calculating Pearson’s correlations between performance in the first and second half of trials.

### Data analyses

We measured the performance of individuals by the proportion of correct responses for each task. We applied Wilcoxon tests followed by Benjamini-Hochberg corrections (for multiple testing) for each task and lemur species to examine whether they performed above chance level. Since no individual solved the social learning task and only one the tool use task, we omitted both tasks from the interspecies comparisons. To analyse whether the three lemur species differed in their performance in the tasks of the PCTB, we used multivariate analysis of variance (MANOVA) with species, sex, rank, age and age:species as between-subject factor and their performance in all tasks as dependent variable. To compare all three species’ performances between the different tasks, we used univariate analysis of variance (ANOVA, for normally distributed data) or Kruskall-Wallis tests followed by *post hoc* analyses (with Bonferroni correction). For significant results, we used an analysis of covariance (ANCOVA) to control for age in these tasks.

Comparisons of performance in tests of the PCTB were conducted between the three lemur species and four haplorhine species (chimpanzees, orangutans, olive baboons, and long-tailed macaques) for which data on individual performance were kindly provided by E. Herrmann and V. Schmitt. On the scale level we applied a MANOVA, followed by ANOVAs or Kruskall-Wallis tests and *post hoc* corrections (Bonferroni) in case of significant results. All statistical analyses were conducted in R version 3.2.2 (R Core Team, Vienna, Austria).

## RESULTS

### Lemurs’ performance in the physical and social domain

In the *physical domain*, the chance level was at 33% in all four tasks of the *scale space*. The three lemur species performed significantly above chance level in the spatial memory and the rotation task (Table 2, Fig. 1). In the object permanence tasks, only ruffed lemurs performed above chance level, while in the control task, all three species performed above chance level (Table 2, Fig. 1). In the *scale quantity*, the three lemur species performed significantly above chance level (50%) in both tasks (Table 2, Fig. 1). In the *scale causality*, the tool use task was successfully solved by only one ring-tailed lemur. However, in the shape and tool properties tasks, all three lemur species performed above chance level (50%; Table 2).

**Table 2.**
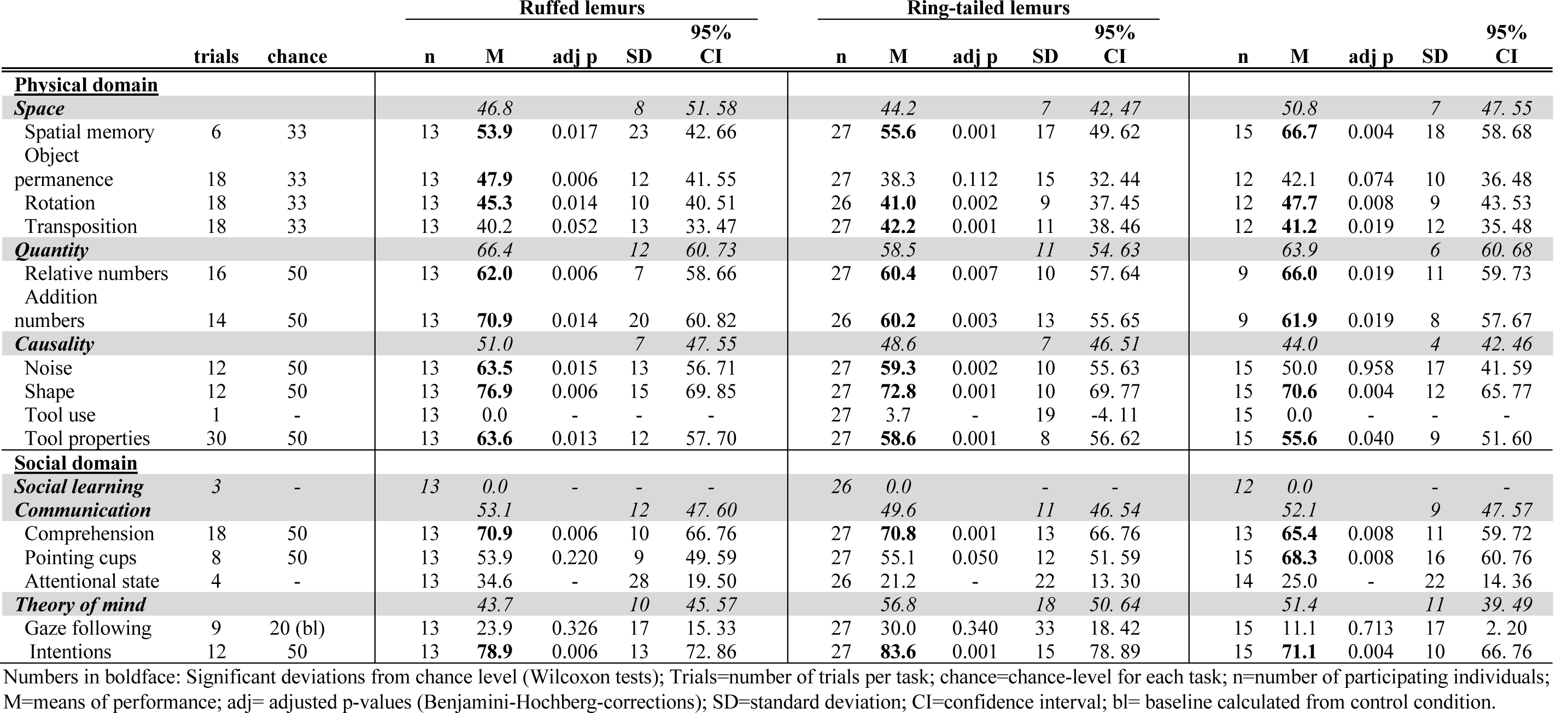
Summary of the mean proportions of correct responses of the three lemur species in all tasks and *scales* of the PCTB.

**Figure 1.**
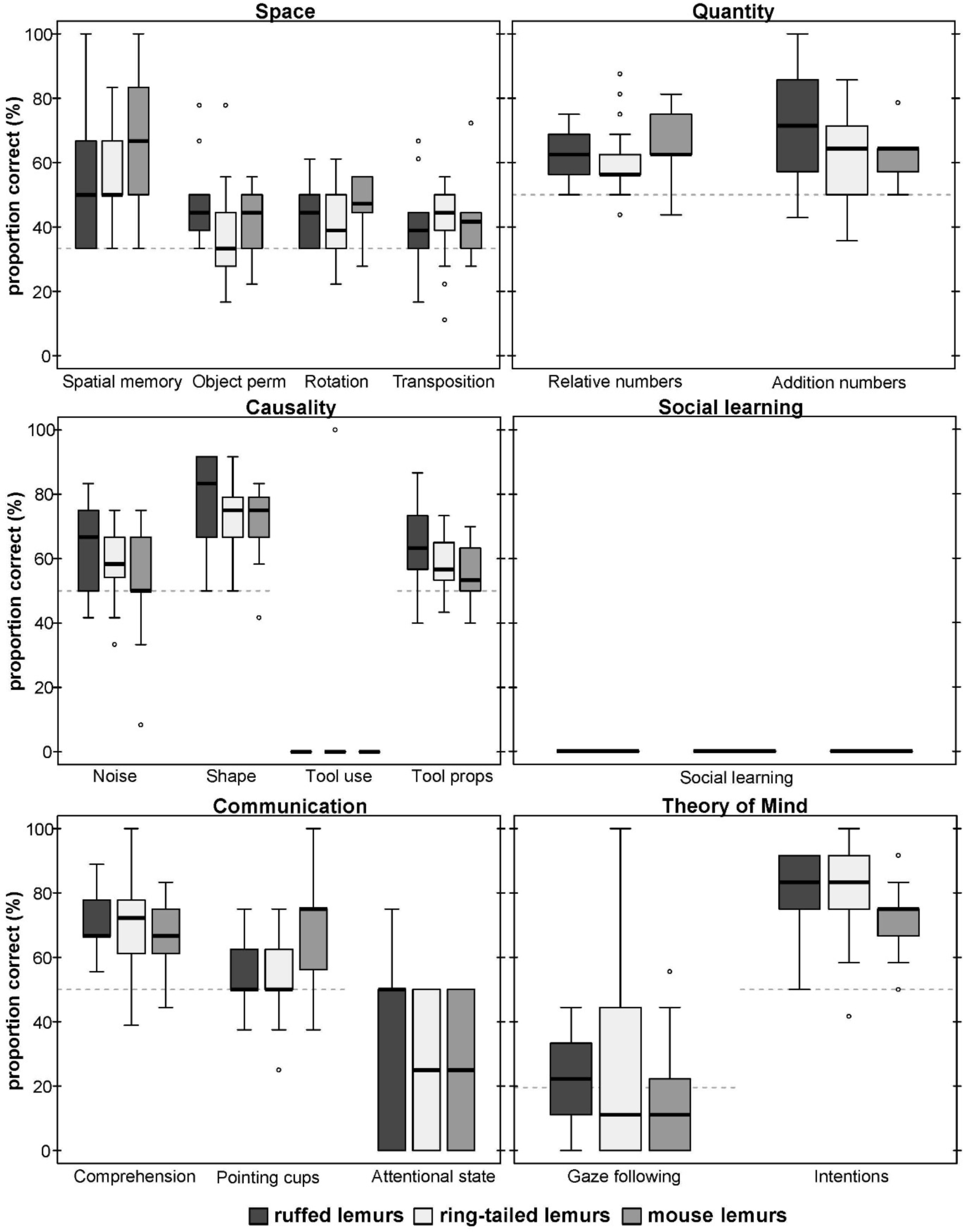
Average performance of the three lemur species in all tasks of the PCTB. Represented are medians (black bars), interquartile ranges (boxes), upper and lower hinges (whiskers), and outliers (circles).

In the *social domain*, no lemur solved the social learning task using a similar technique as demonstrated by a human experimenter (Table 2, Fig. 1). In the *scale communication*, all three lemur species performed significantly above chance level (50%) in the comprehension task, whereas only mouse lemurs performed above chance level (50%) in the pointing cups task. No lemur species performed above chance level in the attentional state task. In the *scale Theory of Mind*, none of the lemur species did follow the gaze of the human experimenter upwards significantly more often than in the control condition in which no cue was given (baseline: 20%; Table 2, Fig. 1). In contrast, all lemur species performed significantly above chance level (50%) in the intentions task (Table 2, Fig. 1).

### Influence of age, sex and rank on performance of the three lemur species

Because the tool use task was solved by only one individual and the social learning task by none, these two tasks were excluded from this comparison. A multivariate analysis of variance of the 14 remaining tasks revealed no differences in the average performance among the three lemur species (MANOVA; Wilk’s Λ=0.498, *F*(19,14)=1.37, p=0.257). Furthermore, average performance was not influenced by sex (Wilk’s Λ=0.461, *F*(19,14)=1.59, p=0.173), rank (Wilk’s Λ=0.273, *F*(38,28)=1.24, p=0.268), age (Wilk’s Λ=0.568, *F*(19,14)=1.03, p=0.466) or age within species (age:species; Wilk’s Λ=0.599, *F*(19,14)=0.91, p=0.566).

### Personality, inhibitory control and learning

The three temperament measures (latency, proximity and duration) of ring-tailed or ruffed lemurs did neither correlate with the performance in the physical domain of the PCTB (Pearson’s correlations, all p>0.05, see Supplemental), nor with the performance of ring-tailed lemurs in the *social domain*. In ruffed lemurs, however, the latency to approach and proximity to a novel stimulus correlated with performance in the social domain (latency to approach: Pearson’s correlation, *r*(11)=0.61, p=0.026; proximity: Pearson’s correlation, *r*(11)=-0.59, p=0.032). No correlation was found between time individuals spent close to the setup (duration) and performance (Pearson’s correlation, *r*(11)=-0.30, p=0.323). Performance in the inhibitory control task did not correlate with performance in the physical and social domain (see Table S4, Supplemental). In addition, we did not find a learning effect in performance between the first and second half of trials within the tasks (Wilcoxon Signed-Rank test: V=806.5, p=0.585).

### Comparison of lemurs and haplorhines in the physical and social domain

The comparison of chimpanzees, orangutans, baboons, macaques, ruffed-, ring-tailed- and mouse lemurs in their overall average performance in the two domains revealed differences among species (Wilk’s Λ=0.383, *F*(406,12)=20.87, p<0.001). Species differed in performance in the *physical domain* (Kruskal-Wallis, *χ*^2^=127.26, df=6, p<0.001; Fig. 2), but not in the *social domain* (Kruskal-Wallis, *χ*^2^=10.25, df=6, p=0.115; Fig. 2). In the *physical domain*, only chimpanzees performed significantly better than ruffed lemurs, and chimpanzees and orangutans outperformed ring-tailed and mouse lemurs (see Table S4, Supplemental).

**Figure 2.**
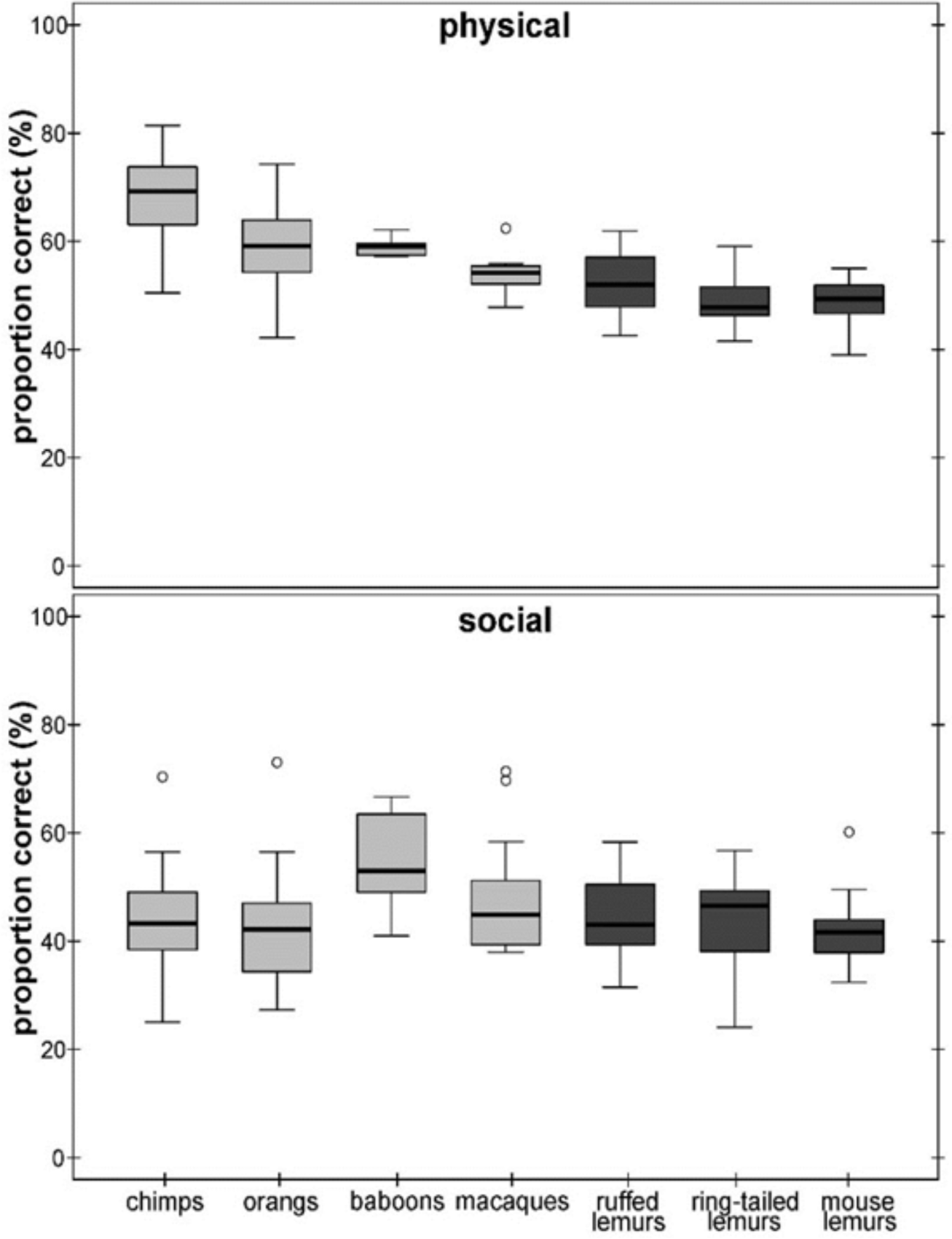
Average performance of apes & monkeys (light grey) and lemurs (dark grey) in the two cognitive domains. Represented are medians (black bars), interquartile ranges (boxes), upper and lower hinges (whiskers), and outliers (circles).

### Comparison of lemurs and haplorhines in the different scales

For a more detailed comparison of all seven species, we conducted a MANOVA including each individuals’ overall performance in all six scales, which revealed significant differences among species (Wilk’s Λ=0.284, *F*(833,36)=7.68, p<0.001). Species differed in all scales except the *scale communication* (ANOVAs or Kruskal-Wallis tests, see Table 3; Fig. 3). In the *scale space*, chimpanzees outperformed all other species, except baboons. Orangutans performed better than ruffed and ring-tailed lemurs, baboons performed better than all three lemur species, and macaques performed similar to all lemur species (Table 4; Fig. 3). In the *scale quantity*, only chimpanzees performed better than ring-tailed lemurs (Table 4; Fig. 3), and in the *scale causality*, chimpanzees outperformed all other species, and orangutans performed better than mouse lemurs (Table 4; Fig. 3). However, this scale was strongly biased by the results of the tool use task, which was only solved by chimpanzees, orangutans and one ring-tailed lemur. Excluding the tool use task from this comparison revealed that only chimpanzees performed better than mouse lemurs (Table 4; Fig. S2, Supplemental).

**Table 3.**
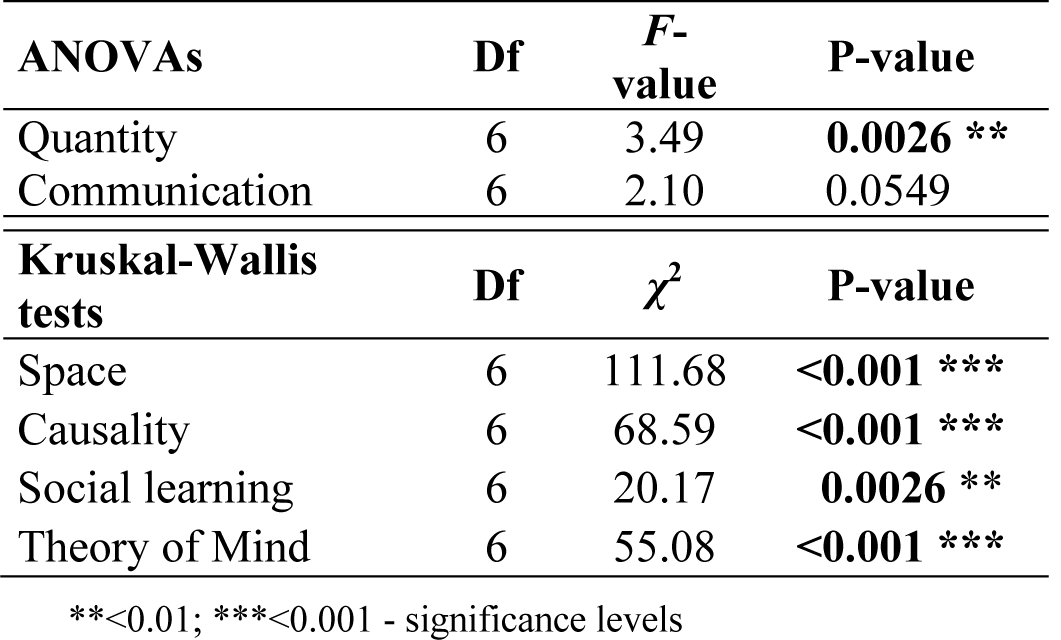
Univariate analyses for the species differences for the six scales.

| **ANOVAs** | **Df** | ***F*-**<br>**value** | **P-value** |
| --- | --- | --- | --- |
| Quantity | 6 | 3.49 | **0.0026** ** |
| Communication | 6 | 2.10 | 0.0549 |
| **Kruskal-Wallis**<br>**tests** | **Df** | **$\chi^2$** | **P-value** |
| Space | 6 | 111.68 | **<0.001** *** |
| Causality | 6 | 68.59 | **<0.001** *** |
| Social learning | 6 | 20.17 | **0.0026** ** |
| Theory of Mind | 6 | 55.08 | **<0.001** *** |

\*\*<0.01; \*\*\*<0.001 - significance levels

**Table 4.** Comparisons of performance among the seven non-human primate species for all six scales of the PCTB. Presented are the results of *post hoc* multiple comparisons (Bonferroni); significant results are in boldface. Causality II: The scale causality without the tools use task.

|  | Space | Quantity | Causality | Causality II | Social learning | Communication | Theory of Mind |
| --- | --- | --- | --- | --- | --- | --- | --- |
| Chimp - Orang | <0.001 | 0.275 | <0.001 | 1 | 1 | 1 | 1 |
| Chimp - Baboon | 1 | 1 | **0.003** | 1 | 1 | 1 | 0.082 |
| Chimp - Macaque | <0.001 | 1 | <0.001 | 1 | 0.699 | 1 | <0.001 |
| Chimp - Ruffed lemur | <0.001 | 1 | <0.001 | 1 | 0.352 | 1 | 0.077 |
| Chimp - Ring-tailed lemur | <0.001 | <0.001 | <0.001 | 1 | **0.025** | 0.29 | <0.001 |
| Chimp - Mouse lemur | <0.001 | 1 | <0.001 | **0.041** | 0.229 | 1 | 1 |
| Orang - Baboon | 1 | 1 | 1 | 1 | 1 | 0.677 | **0.014** |
| Orang - Macaque | 1 | 1 | 0.433 | 1 | 1 | 1 | <0.001 |
| Orang - Ruffed lemur | **0.004** | 1 | 1 | 0.560 | 1 | 1 | **0.009** |
| Orang - Ring-tailed lemur | <0.001 | 1 | 0.643 | 1 | 0.919 | 1 | <0.001 |
| Orang - Mouse lemur | 0.237 | 1 | **0.046** | 0.918 | 1 | 1 | 1 |
| Baboon - Macaque | 0.176 | 1 | 1 | 1 | 1 | 0.591 | 1 |
| Baboon - Ruffed lemur | **0.001** | 1 | 1 | 1 | 1 | 0.653 | 1 |
| Baboon - Ring-tailed lemur | <0.001 | 1 | 1 | 1 | 1 | 0.094 | 1 |
| Baboon - Mouse lemur | **0.023** | 1 | 1 | 1 | 1 | 0.424 | 0.816 |
| Macaque - Ruffed lemur | 1 | 1 | 1 | 1 | 1 | 1 | 1 |
| Macaque - Ring-tailed lemur | 0.074 | 0.307 | 1 | 1 | 1 | 1 | 1 |
| Macaque - Mouse lemur | 1 | 1 | 1 | 1 | 1 | 1 | **0.033** |
| Ruffed lemur - Ring-tailed lemur | 1 | 0.409 | 1 | 1 | 1 | 1 | 1 |
| Ruffed lemur - Mouse lemur | 1 | 1 | 1 | **0.008** | 1 | 1 | 1 |
| Ring-tailed lemur - Mouse lemur | 1 | 1 | 1 | 0.106 | 1 | 1 | **0.036** |

**Figure 3.**
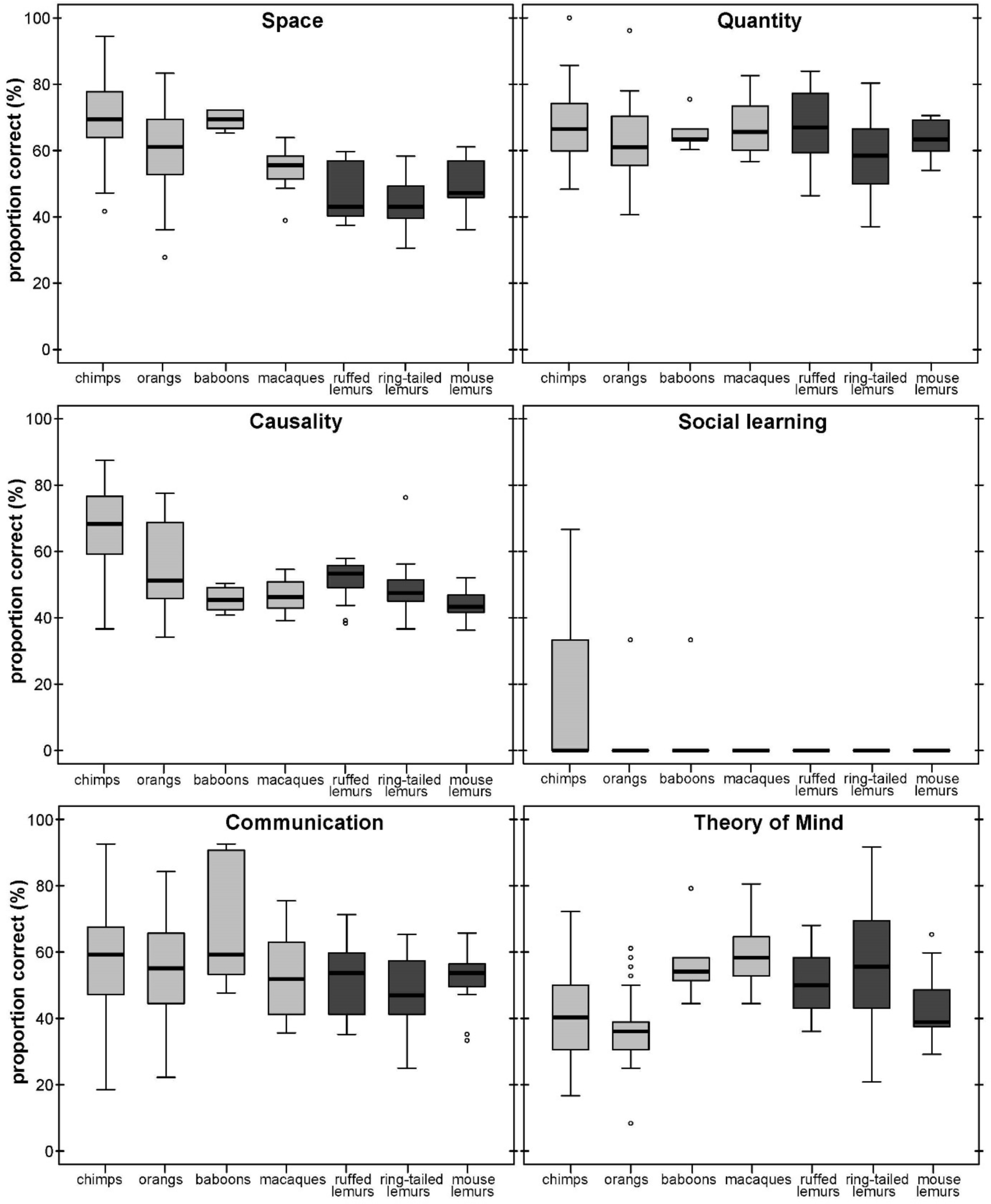
Average performance of apes & monkeys (light grey) and lemurs (dark grey) over the six scales. Represented are medians (black bars), interquartile ranges (boxes), upper and lower hinges (whiskers), and outliers (circles).

In the *social domain*, all species, except great apes, performed poorly in the social learning task, whereas all species performed equally well in the *scale communication* (Table 4; Fig. 3). In the *scale Theory of Mind*, however, chimpanzees performed less good than macaques and ring-tailed lemurs. All other species performed better than orangutans, except mouse lemurs and macaques, and ring-tailed lemurs outperformed mouse lemurs (Table 4; Fig. 3).

## DISCUSSION

In this study, we applied the Primate Cognition Test Battery to three lemur species differing in socioecological traits and brain size and compared their performance with that of four haplorhine species tested in previous studies with the exact same methods. In the *physical domain*, apes and baboons performed better than lemurs in the *space scale*, chimpanzees performed better than ring-tailed lemurs in the *quantity scale* but no differences among species were found in the *causality scale*, after excluding the tool use task. In the *social domain*, lemurs performed at level to apes and monkeys. Most interestingly, in the *Theory of Mind scale*, great apes were outperformed by all other species except mouse lemurs. Since these species differ in relative and absolute brains size (Table 1), with a more than 200-fold difference in brain size between mouse lemurs and orangutans or chimpanzees, our results do not support the notion of a clear-cut link between brain size and cognitive skills, but suggest a more domain-specific distribution of cognitive abilities in primates.

In the *physical domain*, lemurs were outperformed by apes and baboons in the *space scale*. The species with the largest brains (apes and baboons) performed better than all other species, supporting the General intelligence hypothesis. These findings are in line with an earlier study showing that apes and monkeys differ in their ability to track object displacements (Amici et al. 2010). Spatial understanding is also important to remember food resources or to track conspecifics (Dunbar and Shultz 2017), and species (chimpanzees, orangutans, baboons) having a larger dietary breadth performed better in these tasks, but the species with the highest amount of fruits in the diet (ruffed lemurs) did not perform better than other species, providing only partial support for the Ecological intelligence hypothesis. There was no clear pattern between group size and performance in the *space scale*, providing no support for the Social intelligence hypothesis.

In the *quantity scale*, only chimpanzees performed better than ring-tailed lemurs, and all other species performed similarly, indicating that a certain level of numerical understanding appears to be a basal cognitive trait of all primates. These results support earlier studies indicating that lemurs do not differ from haplorhine primates in numerosities and simple arithmetic operations (Jones and Brannon 2012; Merritt et al. 2011; Santos et al. 2005a). Since a comparable numerical understanding as tested in the PCTB has also been reported for various taxa outside the primate order, including fish and insects (e.g. Agrillo et al. 2012; Chittka and Geiger 1995; Pahl et al. 2013; but see Krasheninnikova et al. 2019), a basal numerical understanding may be present in many animals.

In the *causality scale*, lemurs performed as well as both monkey species, but all monkeys and lemurs were outperformed by chimpanzees, who excelled in the tool use task. Even natural tool users, such as orangutans and long-tailed macaques (Brotcorne et al. 2017; van Schaik et al. 2003), hardly solved this task (Schmitt et al. 2012). It required the ability to use a stick to rake a food reward into reach, which might have been too challenging for species exhibiting either a medium (baboons, macaques) or low (lemurs) level of precision grip (Torigoe 1985). Although long-tailed macaques use stone tools to crack open nuts or mussels, they do so mainly by applying force rather than using fine-motor skills (Gumert and Malaivijitnond 2012). Thus, the tool use task appears unsuitable for a fair interspecific comparison. Excluding this task from the *quantity scale* resulted in a rather similar overall average performance of all species. Interestingly, lemurs that have never been observed to use tools in the wild (Fichtel and Kappeler 2010; Kittler et al. 2015, 2018), appeared to exhibit an understanding for the necessary functional properties of pulling tools (Santos et al. 2005b; Kittler et al. 2018). Hence, except for the *space scale* we did not find systematic species differences in performance, challenging the notion that there is a domain-general distinction between haplorhines and strepsirrhines (Deaner et al. 2006). Our results instead suggest the existence of domain-specific cognitive differences.

In the social domain, species differences were less pronounced, and lemurs’ overall performance in the *Theory of Mind scale* was equal to that of monkeys and even superior to that of apes. In the *social learning scale* neither lemurs, nor baboons or long-tailed macaques solved the task. However, long-tailed macaques exhibit cultural variation in stone handling techniques in the wild, indicating that they are able to learn socially (Brotcorne et al. 2017). The ability to learn socially has also been reported in ring-tailed and ruffed lemurs (e.g. Kappeler 1987; Kendal et al. 2010; O’Mara and Hickey 2012; Stoinski et al. 2011), but remains unstudied in mouse lemurs. Since individuals had to learn in this task from a human demonstrator, the phylogenetic distance between species and the demonstrator might have influenced learning abilities, because great apes performed better than Old World monkeys and lemurs (Schmitt et al. 2012). Hence, it remains an open question whether monkeys and lemurs would perform better when tested with a conspecific demonstrator. Moreover, the task required the ability to shake a transparent tube or to insert a stick into the tube, which might have been too difficult for species with limited dexterity (Torigoe 1985). Therefore, a social learning task adapted to manipulative skills of Old World monkeys and lemurs (Schnoell and Fichtel 2012) might be more informative in future studies.

In the *communication scale*, all species performed equally well, suggesting that all species can make use of socio-visual cues given by others. This result is in line with those of several other studies showing the ability to use social-visual cues presented by a human demonstrator in object-choice experiments in birds (Schmidt et al. 2011), aquatic mammals (sea lions: Malassis and Delfour 2015; dolphins: Tschudin et al. 2001), domestic animals (dogs: Kaminski et al. 2005; Miklósi et al. 1998; pigs: Nawroth et al. 2016; goats: Wallis et al. 2015), as well as other primates (Anderson et al. 1995; Itakura 1996).

In contrast, unexpected species differences emerged in the *Theory of Mind scale*, with great apes performing inferior to both monkeys and lemurs. This difference was mainly due to better performance of monkeys and lemurs in the intentions task, but not in the gaze following task. In the gaze following task all lemurs performed below chance level, although it has been shown that ring-tailed lemurs follow the gaze of conspecifics (Shepherd and Platt 2008) and that they use human head orientation as a cue for gaze orientation in a food choice paradigm (Botting et al. 2011, Sandel et al. 2011), questioning the validity of these gaze following tasks. In the intention task, a human observer tried to reach a cup with a hidden reward repeatedly with the hand. Monkeys and lemurs might have performed better than apes because they may have solved the task by using spatial associations between the repeated hand movements and the cup or by understanding the hand movements as a local enhancement (Shettleworth 2010; Schmitt et al. 2012). Still, it remains puzzling why chimpanzees and orangutans did not use the hand movement as a cue for the location of the hidden reward. Even more so because a comparative study of Theory of Mind compatible learning styles in a simple dyadic game between seven primate species, including chimpanzees and ring-tailed lemurs, and a competitive human experimenter revealed that test performance was positively correlated with brain volume but not with social group size, suggesting that Theory of Mind is mostly determined by general cognitive capacity (Devaine et al. 2017). Hence, additional social cognitive tests are required to obtain a better understanding of the relationship between brain size and cognitive performance in the social domain.

Altogether, average species performances were generally not as different as it might have been expected in view of the various hypotheses on the evolution of cognitive abilities. Except for the scale space, the overall comparison does not provide support for the *General intelligence hypothesis*, since variation in brain size cannot explain the observed results. Similarly, performances of the seven species did not reflect any clear patterns concerning their feeding ecology, i.e. the percentage of fruit in the diet or dietary breadth, except for the *space scale* (see Table 1); hence, these results do not provide support for the *Ecological intelligence hypothesis*. Moreover, our results do not provide support for the *Social intelligence hypothesis* because lemurs, and especially the solitary mouse lemurs, should have performed inferior compared to the haplorhine species (Dunbar and Shultz 2017).

Earlier comparative studies among primates linking performance in a range of comparable cognitive tests in the physical or social domain revealed a link between performance in these tasks and brain size (Deaner et al. 2006, 2007; Reader and Laland 2002; Reader et al. 2011). However, studies using the exact same experimental set up revealed contradictory results. Two studies addressing only one cognitive ability revealed a positive relationship between brain size and performance in inhibitory control or Theory of Mind tests (Maclean et al. 2014; Devaine et al. 2017), but all other studies applying various tests on inhibitory control and spatial memory (Amici et al. 2008, 2010, 2012) or tasks of the Primate Cognition Test Battery (Herrmann et al. 2007; Schmitt et al. 2012; this study), found no clear-cut relationship between brain size and cognitive performance.

Even though lemurs performed at level with monkeys and great apes in many of these experiments, we do not suggest that their cognitive abilities are *per se* on par with those of larger-brained primates. In the physical domain, species differences emerged only in the space scale, supporting the General intelligence hypothesis. However, no systematic species differences were found in the quantity or causality scales, which address rather basal cognitive abilities, which might not be variable enough to reveal actual differences between species. Indeed, some fish and insects possess similar basal cognitive skills in the physical domain (Fuss et al. 2014; Loukola et al. 2017; Schluessel et al. 2015). In the social domain, the social learning task was not suitable for all species, and individuals might have recruited other abilities to solve the problems, as discussed for the intention task above.

Many tests of the PCTB were based on two-or three-choice paradigms in which the costs for choosing correctly were rather low, because the probability to receive a reward was either 50% or 33%, a random choice strategy might have been still relatively profitable. For example, performance in a memory task increased in common marmosets (*Callithrix jacchus*) and common squirrel monkeys (*Saimiri sciureus*) from a two-choice task to a nine-choice task, in which the probability of success was lowered from 50% to 11%, making a wrong choice more costly appeared to favour an appropriate learning strategy over a random choice strategy (Schubiger et al. 2016). The application of a random choice strategy may also explain why four parrot species that have been tested with the PCTB, may have failed to solve the tasks, besides morphological differences in performing the tasks (Krasheninnikova et al. 2019).

In addition, the PCTB was designed to examine the spontaneous ability to solve the tasks, and not to examine how long individuals need to learn the task. Hence, a test battery that continued testing until individuals reached a certain criterion (e.g. 80 % correct responses) or detailed analyses of applied learning strategies as in Devaine et al. (2017) may allow to compare not only species differences in their spontaneous ability to solve the task, but also species-specific learning curves as well as learning strategies, which might reveal more informative differences.

To conclude, our study generated the first systematic results on cognitive abilities in lemurs, and the comparison with haplorhines suggests that in many aspects of the physical and social domain, the average performance in these tests of members of these two lineages do not differ substantially from each other. These results reject the notion of a direct correlation between brain size and cognitive abilities and question assumptions of domain-general cognitive skills in primates. Overall, our results strengthen the view that when comparing cognitive abilities among species, it is of vital importance to include a diverse set of tests from both cognitive domains which are applicable to a diverse range of species and taxa (Auersperg et al. 2011, 2013; Burkart et al. 2016; MacLean et al. 2012; Schmitt et al. 2012) and to carefully consider the external validity of the specific tests (Krasheninnikova et al. 2019).

## ACKNOWLEDGEMENTS

We are grateful towards Silvio Dietzel and the “Erlebnispark Affenwald” for permission to work with their lemurs. We would also like to thank Esther Herrmann and Vanessa Schmitt for sharing the PCTB performance data of the great apes and monkeys with us. Furthermore, we are grateful to Ulrike Walbaum, Anna Zango Palau, Luise Zieba and Lluis Socias Martinez for helping with the experiments and inter-observer coding the videos. Thanks to Sarah Hartung, Henry Benseler and Ramona Lenzner-Pollmann for taking care of the animals. This study was supported by the DFG (awarded to CF: FI 929/8-1).

## AUTHORS’ CONTRIBUTIONS

C.F. and K.D. have a shared first-authorship, they conceived the study and analysed the data. K.D. conducted the experiments. C.F., K.D. and P.M.K. wrote the manuscript. All authors gave final approval for publication.

## COMPETING INTERESTS

We have no competing interests.

## Notes

### Competing Interest Statement

The authors have declared no competing interest.

