## Supplemental for "The lemur baseline: How lemurs compare to monkeys and apes in the Primate Cognition Test Battery"

### 1. Detailed Methodology of the Primate Cognition Test Battery (*PCTB*)

Experimental setups were adopted from Herrmann et al. (2007) and Schmitt et al. (2012). As suggested by Schmitt et al. (2012) we doubled the number of trials for all object-choice tasks from 3 to 6 (see Table S1) to include all possible locations and combinations. In addition, some of the original tasks were extended by using control conditions and the quantity combinations in experiments 5 and 6 were adopted from Schmitt et al. (2012). Otherwise the experimental setups are the same in the PCTB by Hermann et al. (2007). To avoid confusion, we used a similar wording to describe the tasks. The size of the items and objects used was adjusted to make them operable for lemurs, especially for the small mouse lemurs. All experiments were conducted by the same experimenter (E1), except those for which a second person (E2) was required (1.2.3, 1.2.4 and 1.2.6). As second person (E2), two different persons assisted, but the same person always assisted in one task for all trials and individuals.

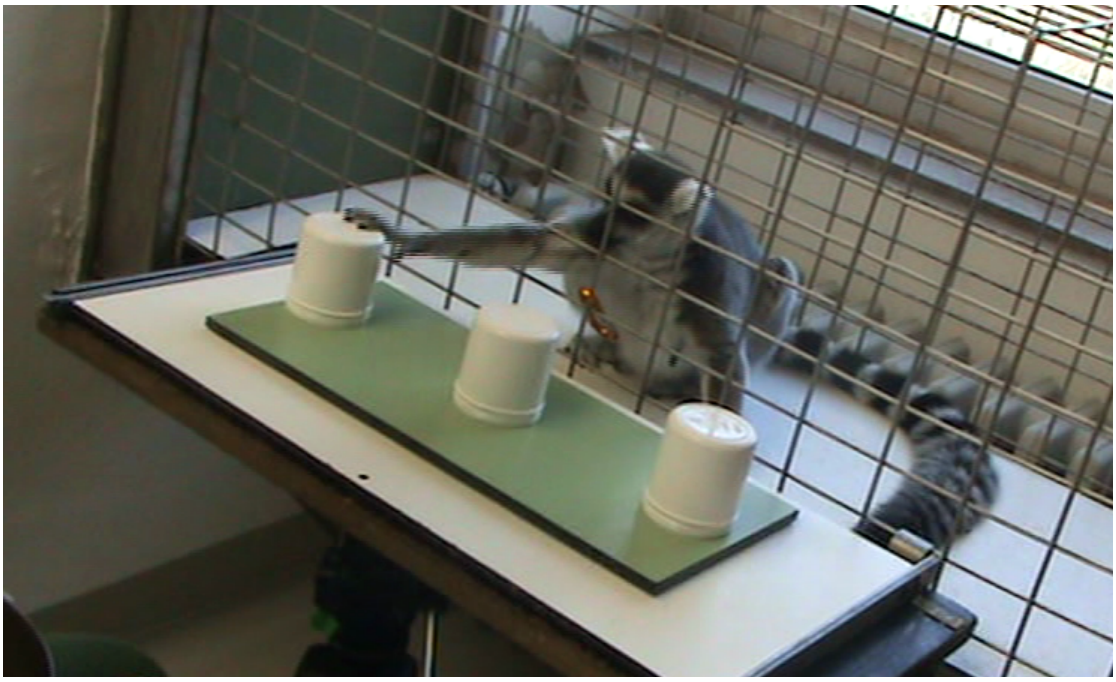

**Figure S1** Basic experimental setup of the PCTB (depicted here experiment 3, *Rotation*). The table with the sliding board on top is attached to the mesh of the subjects' cage and the subject is positioned in the centre of the setup using a carabiner (they have been previously trained to stay put wherever the carabiner is positioned). In this example, after watching the placement of the reward and the subsequent rotational movement, the sliding board was pushed towards the individual to choose between the three cups.

**Table S1** Summary of the PCTB and the number of trials per task and individuals per species and task.

| | scale | task | trials | *N tested ruffed lemurs* | *N tested ring-tailed lemurs* | *N tested mouse lemurs* |
|---|---|---|---|---|---|---|
| ***physical*** | space | **2.1.1** ***Spatial memory*** | **6** | 13 | 27 | 16 |
| | | **2.1.2** ***Object permanence*** | **24** | 13 | 27 | 12 |
| | | a) Single displacement | 6 | | | |
| | | b) Double-adjacent displacement | 6 | | | |
| | | c) Double non-adjacent displacement | 6 | | | |
| | | d) Single displacement touch | 6 | | | |
| | | **2.1.3** ***Rotation*** | **18** | 13 | 26 | 12 |
| | | a) 360° | 6 | | | |
| | | b) 180° middle | 6 | | | |
| | | c) 180° side | 6 | | | |
| | | **2.1.4** ***Transposition*** | **18** | 13 | 27 | 12 |
| | | a) Single | 6 | | | |
| | | b) Double unbaited | 6 | | | |
| | | c) Double baited | 6 | | | |
| | quantities | **2.1.5** ***Relative numbers*** | **16** | 13 | 27 | 9 |
| | | **2.1.6** ***Addition numbers*** | **14** | 13 | 26 | 9 |
| | causality | **2.1.7** ***Noise*** | **12** | 13 | 27 | 15 |
| | | a) Noise full | 6 | | | |
| | | b) Noise empty | 6 | | | |
| | | **2.1.8** ***Shape*** | **12** | 13 | 27 | 15 |
| | | a) Board | 6 | | | |
| | | b) Cloth | 6 | | | |
| | | **2.1.9** ***Tool use*** | **1** | 13 | 27 | 16 |
| | | **2.1.10** ***Tool properties*** | **30** | 13 | 27 | 15 |
| | | a) Side | 6 | | | |
| | | b) Bridge | 6 | | | |
| | | c) Ripped | 6 | | | |
| | | d) Broken wool | 6 | | | |
| | | e) Tray circle | 6 | | | |
| ***social*** | social learning | **2.2.1** ***Social learning*** | **4** | 13 | 26 | 15 |
| | | a) Paper tube | 1 | | | |
| | | b) Banana tube | 1 | | | |
| | | c) Stick tube | 1 | | | |
| | communication | **2.2.2** ***Comprehension*** | **42** | 13 | 27 | 13 |
| | | a) Head & eyes | 18 | | | |
| | | b) Head, eyes & paw | 18 | | | |
| | | c) Marker | 6 | | | |
| | | **2.2.3** ***Pointing cups*** | **8** | 13 | 27 | 15 |
| | | **2.2.4** ***Attentional state*** | **4** | 13 | 26 | 15 |
| | | a) Away | 1 | | | |
| | | b) Towards | 1 | | | |
| | | c) Away body-facing | 1 | | | |
| | | d) Towards body-facing | 1 | | | |
| | theory of mind | **2.2.5** ***Gaze following*** | **9** | 13 | 27 | 16 |
| | | a) Head & eyes | 3 | | | |
| | | b) Back | 3 | | | |
| | | c) Eyes | 3 | | | |
| | | **2.2.6** ***Intentions*** | **12** | 13 | 27 | 15 |
| | | a) Trying | 6 | | | |
| | | b) Reaching | 6 | | | |

### 1.1 Tasks of the physical domain

#### 1.1.1 Spatial memory

On the experimental sliding board three cups were placed in a row. While the individual was watching, two rewards were first presented and then openly placed under two of three cups. After the board was moved towards the individual the individual had to choose between the three cups and point at the chosen one. The individual could choose two times consecutively, but if it chose the cup without reward first, no further choices were allowed. Individuals had to choose both cups correctly to count as a correct response.

#### 1.1.2 Object permanence

Again, three cups were placed in a row. A smaller fourth cup (placed in the beginning on the far left or right side of the board) was used for displacing the reward into one of these three cups. Therefore, while the individual was watching, a reward was placed under the fourth cup and afterwards four different displacement-scenarios were conducted:

a) *Single displacement*: The fourth cup, including the reward, was moved under one of the three big cups without touching the other two cups.
b) *Double adjacent displacement*: The fourth cup was moved consecutively under two adjacent cups and the reward was left under one of these cups without touching the third.
c) *Double non-adjacent displacement:* The fourth cup was moved under the two outer cups and the reward was left under one of these cups without touching the cup in the centre.
d) *Single displacement touch*: The fourth cup was moved under one of the three cups and the reward was left there. E1 touched the other two cups in order to find out whether the individuals simply chose the cup touched last by E1 or indeed followed the small cup to the last location it was moved to.

After these displacements, the empty fourth cup was shown to the individual and the board was moved towards it. The individual was now allowed to make one choice for the single displacement and two consecutive choices for the double displacements. If it chose a cup that had not been part of the displacement-scenario no further choices were allowed. Individuals had to choose the reward-cup as first choice to count as a correct response.

#### 1.1.3 Rotation

A movable tray is put on top of the board with three cups placed on it in a row. While the individual was watching, a reward was first presented and then openly placed under one of the three cups. The tray and hence the cups were then rotated in three different spatial scenarios:

a) *360°:* The reward was placed under one of the outer cups and the tray was rotated 360° in clockwise (or counter clockwise) direction. Hence, the reward was in the end again in the same position as before the rotation.
b) *180° middle:* The reward was placed under the cup in the centre and the tray was rotated 180° in clockwise (or counter-clockwise) direction. Hence, the reward was in the end still in the same position as before the rotation.
c) *180° side:* The reward was placed under one of the outer cups and the tray was rotated 180° in clockwise (or counter-clockwise) direction. Hence, the reward was in the end in the opposite position as before the rotation.

After the rotations, the board was moved towards the individual and it could choose a cup once. Individuals had to choose the reward-cup correctly to count as a correct response.

#### 1.1.4 Transposition

Again, three cups were placed in a row and while the individual was watching a reward was first presented and then openly placed under one of the cups. The cups were then transpositioned in three different spatial scenarios:

a) *Single transposition:* The position of the reward-cup was switched with one of the empty cups without touching the third cup.
b) *Double unbaited transposition:* The position of the reward-cup was switched with one of the empty cups and afterwards the positions of the two empty cups were switched.
c) *Double baited transposition:* The position of the reward-cup was switched with one of the empty cups and afterwards again switched with the other empty cup.

After the transpositions, the individual could make a choice once and only the reward-cup being the first choice counted as a correct response.

#### 1.1.5 Relative numbers

Two plastic plates were placed on the testing board and then hid from the view of the individual using an occluder. Both plates were then baited with different amounts of equally sized reward pieces, covered with lids and placed in the middle of the board. After removing the occluder the lids of both plates were simultaneously lifted and hence the individual could see the amounts of reward pieces in each plate for about 5 seconds. Then the plates were moved to the sides of the board, one right and one left, and the individual could make its choice. Each of the following pairs of numbers of reward pieces was trialled once per individual[2] (the order of presentation was randomized):

*1:0 II 1:2 II 1:3 II 1:4 II 1:5 II 2:3 II 2:4 II 2:5 II 2:6 II 3:4 II 3:5 II 3:6 II 3:7 II 4:6 II 4:7 II 4:8*

(Additional four control conditions *1:1 II 2:2 II 3:3* and *4:4* were tested to monitor any possible side biases, e.g. choosing the same side in every trial.)
The individual had to choose the larger quantity first to count as a correct response.

#### 1.1.6 Addition numbers

Hidden behind the occluder, three plastic plates were baited with different amounts of reward pieces and then covered with lids and placed in the middle of the board. The occluder was removed, the lids of the outer plates were lifted simultaneously, and the individual could see them for about 5 seconds. Then they were covered again, and the lid of the middle plate was uncovered, allowing the individual to see its amount of reward pieces for 5 seconds. Afterwards the contents of the middle plate were transferred into one of the outer plates, with the individual being able to watch the transfer but not the content of the side plates. The empty middle plate was removed from the board and the individual could make its choice between the two covered outer plates. Each of the following pairs of reward pieces is trialled once per individual (the order in which they are presented is randomized):
*1:0 + 3:0 = 4:0 II 6:1 + 0:2 = 6:3 II 2:1 + 2:0 = 4:1 II 4:3 + 2:0 = 6:3 II 4:0 + 0:1 = 4:1 II 2:1 + 0:2 = 2:3 and 4:3 + 0:2 = 4:5*
(Each combination was presented with the resulting higher number being once on the left and once on the right side, resulting in 14 trials in total.) The individual had to choose the larger quantity first to count as a correct response.

#### 1.1.7 Noise

Behind the occluder a reward was hidden in one of two opaque cups. After the occluder was removed, the cups were manipulated in the two following ways while the individual was watching, and it had to choose the reward cup first to count as a correct response:

a) *Noise full:* The reward cup was shaken three times, letting the food rattle inside and the empty one was simply lifted once without shaking (which cup first was randomized).
b) *Noise empty:* The empty cup was shaken three times, producing no sound and the baited cup was simply lifted once without shaking (which cup first was randomized).

#### 1.1.8 Shape

Behind the occluder a reward was hidden beneath one of two identical pieces of plastic board or cloth, thereby changing the appearance of the baited piece. After removing the occluder the individuals were presented with two different situations and they could choose once between the two possibilities. The individual had to choose the reward board or cloth first to count as a correct response.

a) *Board:* The reward was hidden underneath one of two plastic boards (sized 15x10 cm; 4x3 cm for mouse lemurs). The reward plastic board was not lying flat on the surface but inclined a bit.
b) *Cloth:* The reward was hidden underneath one of two pieces of cloth (sized 15x10 cm; 4x3 cm for mouse lemurs). A visible bump in the cloth was made by the reward instead of remaining flat on the surface.

#### 1.1.9 Tool use

A reward was placed on the board out of reach of the individual (about 25 cm; 8 cm for mouse lemurs). Because the reward itself was out of reach for the individual it could only gain the food item by manipulating the tool, in this case a simple wooden stick (length 30 cm; 10 cm for mouse lemurs) that was provided to the individual. It had to retrieve the reward using the tool within two minutes; otherwise the attempt was not counted as a correct response.

#### 1.1.10 Tool properties

Behind the occluder two different tool setups, one intact and effectively functioning to gain the food reward and the other not, were placed on the sliding board. The individual could choose a tool once by pulling it and the first choice had to be the functioning tool to count as a correct response. Five different tool setups and objects were used:

*a)* *Side:* Two identical pieces of cloth (sized 15x10 cm; 4x3 cm for mouse lemurs) were placed next to each other on the board. On top of one piece a reward was placed and for the other piece it was placed directly next to the cloth, making it the ineffective tool. The individual could only gain the reward placed on top of the cloth by pulling at it.
*b)* *Bridge:* Again, two identical pieces of cloth (see above) were placed on the board, but this time two identical plastic bridges were placed over each of their far ends. For the ineffective tool, the reward was placed on top of the bridge and for the other underneath it. Hence the individual could obtain the reward by pulling the cloth.
*c)* *Ripped:* Two pieces of cloth were again used, but only one of them intact the other was ripped apart in the middle. The two broken pieces were placed on the board with a gap of 1 cm in between, making it visually obvious that they were not connected. It was important that the intact piece of cloth (sized 15x10 cm; 5x3 cm for mouse lemurs) was equally sized as the ripped pieces including the gap (2 smaller pieces sized 7x10 cm; 2x3 cm for mouse lemurs). For both cloths, the reward was placed on top of the far end, hence for the ripped cloth on the unreachable piece, making it ineffective. The individual could choose one cloth and obtain the reward by pulling at it.

d) *Broken wool:* This task was basically identical to the previous one, except that pieces of wool string were used instead of cloth. The rewards were tied to the far ends of the wool pieces, making the broken one ineffective. The individual needed to pull at the intact string in order to gain a reward.

e) *Tray circle:* Two small plastic trays (sized 6x6.5 cm; 2x2.5 cm for mouse lemurs) were placed on the board. One of them had a round hole cut in the middle (Ø 3 cm; 1 cm) and the other a u-shaped hole cut from out of its back. A reward was placed in the middle of each of the holes with the round one surrounding it effectively and the u-shaped one not holding it when pulled towards the individual. Using a string attached to the trays the individual was then allowed to pull at one of them to obtain the reward. Only the tray with the round hole would work effectively as it would push the reward towards the cage.

### 1.2 Tasks of the social domain

#### 1.2.1 Social learning

In the three different treatments of this task there was always a piece of reward stuck inside a plastic tube and E1 demonstrated the solution of this problem to the individuals once. The observing individual was then given two minutes to solve the problem on its own. A trial counted as correct only if the individual obtained the reward successfully by using a method highly similar to the previously demonstrated one.

a) *Paper tube:* A reward was placed inside a 10 cm long transparent plastic tube with a piece of paper attached over both ends. E1 demonstrated how to open the tube: First E1 poked her finger through the paper on one end and then wiggled her finger in the tube to rip the paper further, making the hole in the paper larger (i.e. as opposed to using her mouth or hands to tear the paper off the tube). Finally, E1 tilted the tube in order to let the reward fall in her hand. After the demonstration, an identical tube was handed to the individual.

b) *Banana tube:* A small slice of banana was placed in the centre of a transparent plastic tube (15 cm) and a specific force had to be applied to get the reward out of the tube. E1 demonstrated how to get the reward by banging one end of the tube on the table (as opposed to shaking it forcefully). An identical tube with banana inside was afterwards handed to the individual.

c) *Stick tube:* An opaque plastic tube with caps on each end was baited with a reward. One of the caps had a hole in it but was glued to the tube, whereas the other cap had no hole but could be removed. E1 demonstrated how to open the tube: First E1 inserted a stick through the cap with a hole, and then she pushed the stick through the hole which forced the cap on the other end to fall off. After the successful demonstration, an identical grey tube was handed to the individual.

#### 1.2.2 Comprehension

Behind the occluder a reward was hidden under one of two cups placed on the board in a row. After the occluder was removed E1 indicated the rewards' hidden location through three different possible pointing cues:

a) *Look (Head & eyes):* E1 alternated her gaze three times between the individual and the baited cup while calling the individuals' name and afterwards continuously looked towards the cup until the individual chose.
b) *Point (Head, eyes & hand):* E1 alternated her gaze three times between the individual and the baited cup while calling the individuals' name and continuously looked towards the cup and additionally pointed at it with the extended index finger of her cross-lateral hand until the individual chose.
c) *Marker:* E1 held an iconic photo marker, which depicted the reward (banana pieces), in her hand and alternated her gaze three times between the photo and the individual while calling the individuals' name. Then E1 placed the photo on top of the baited cup.

   After the board was moved towards the individual it could choose between both cups. It had to choose the reward-cup first to count as a correct response.

#### 1.2.3 Pointing cups

Two identical cups were placed at the far ends of the sliding board. The individual was directed to a starting point in the middle of the board and E2 (a second experimenter) entered the testing area, placed a reward under one of the two cups while the individual was watching and left again. After E2 left, E1 entered the area and centred the individual again on its starting point using a piece of food. Then E1 stood in the middle between both cups and waited for the individual to choose one of the cups. Given the individual chose the reward cup correctly within 60 seconds E1 offered it the reward and a correct response was scored.

#### 1.2.4 Attentional state

In the beginning a second experimenter (E2) entered and placed a reward in front of the cage but out of reach of the individual, randomly varied either on its right or left side, and left the room again. Afterwards E1 entered but stood at the end of the room opposite of the reward and thus did not notice the reward on the floor. The attentional state of E1 varied in the different trials by looking in 4 different directions:

a) *Away:* E1 turned around and looked away from the reward. The individual had to approach from her front to gain her attention. If the individual did so within 20 sec, E1 turned around and waited for the individual to direct her attention to the reward by moving back to it. If the individual indicated the rewards' location within 20 sec, E1 handed it to the individual.

b) *Towards:* E1 looked towards the reward and waited for the individual to approach the reward and direct her attention towards it within 20 sec. If the individual indicated the rewards' location within 20 sec, E1 handed it to the individual.
c) *Away Body-facing:* This trial was identical to "*Away*", except that E1s' body faced toward the reward and only her face was turned away.
d) *Towards Body-away:* This trial was identical to "*Towards*", except that E1s' body was turned away and only her face was directed towards the reward.

Indicating the location of the hidden food item was possible by pointing to the rewards' location if it was in view of E1 or by first moving into E1s' view (i.e. gaining her attention) and then pointing to the location. To count as a correct response, the individual had to successfully gain the reward by indicating its location to E1, otherwise E1 left the room and E2 entered again and removed the reward.

#### 1.2.5 Gaze following

By calling and presenting a reward the attention of the individual was attracted and once the individual looked at E1 one of three different communicative gaze cues (implemented on different days to minimize any kind of habituation) was performed by orienting in the corresponding direction:

a) *Head & eyes:* E1 called the individuals' name and showed a reward. Then she hid the food in her hand, which remained in front of her body. Afterwards she looked up with both her head and eyes for ~10 sec.
b) *Back*: E1 sat with her back facing the individual. She called the individuals' name and showed a reward. Then she hid the food in her hand, which remained in front of her body. Afterwards she looked up at the ceiling for ~10 sec. Within the ~10 sec she looked back over her shoulder at the individual three times to ensure the individuals' attention. If it was not paying attention when E1 looked the second time, the trial was repeated.
c) *Eyes:* E1 called the individuals' name and showed a reward. Then she hid the food in her hand, which remained in front of her body. Afterwards she looked up at the ceiling for ~10 sec while her face was still facing the individual.

To count as a correct response the individual had to follow the gaze of the conspecific during the first 10 seconds after the E1 changed the gaze direction.

#### 1.2.6 Intentions

Behind the occluder a reward was hidden by E1 in one of two closed metal tins placed on the board in a row. The occluder was removed and the tins were manipulated by E2 as follows:

a) *Trying:* E2 tries in vain to open the reward tin by removing the lid while looking at the tin.

b) *Reaching:* E2 tries in vain to reach for the reward tin by extending the equilateral arm and looking at the tin, but a Plexiglas barrier blocks the access to the tin. The cue is given continuously until the individual indicates its choice.

About 3 seconds after each demonstration E1 approaches again and moves it towards the individual, allowing it to make a choice. The reward tin had to be chosen first to count as a correct response.

### 2. The Personality Study

As temperament or personality of individuals can influence performance in problem solving tasks (Hare and Tomasello 2005), study individuals were tested with respect to their reaction to 29 different items (novel objects, persons, foods; Herrmann et al. 2007; Schmitt et al. 2012). The testing situations varied depending on 1) the nature of the different items presented (humans, objects or food pieces), 2) whether the items were presented in combination or alone (e.g. non-familiar human moving a novel object), and 3) the level of activity of the items that took place during their presentation (e.g. novel object moving, also see Table S2).

The items were presented by a second, unfamiliar experimenter (E2) sitting in front of the cage (except the very first test, in which the familiar human experimenter E1 is presented to the individual). The individual was then directed to a starting point offering food and the stimuli ware each presented for 30 seconds. Each individual participated in one session per day on three consecutive days with the same order of stimuli (see Table S2). The first day all the items were presented and placed on the board by E2 and the individual was only allowed to view them (visible). Additionally, two non-social trials were also run during which the individual could either view the empty board alone or a bright red spot was placed on the board before E2 left the area. During the sessions of the second day (movement) E2 moved the different items from left to right over the board and on the third day (touch) the items were put close to the cage allowing the individual to potentially touch them if they wanted to.

All experiments were videotaped. Each individual's degree of anxiety and/or disinterest as a response to the different testing situations was scored from the video recordings. Therefore, the time it took the individual to approach the new item (latency) or whether they did it at all was noted and also whether they tried to touch it or not. In addition, the time the individual spent near the item (duration) and also how close it approached (proximity) was noted. Overall, this part of the study allows controlling for temperamental factors (anxiety-boldness/interest-disinterest) influencing cognitive abilities.

**Table S2** Summary of the items and methods used in the *Personality Study*.

| | category | item | description |
|---|---|---|---|
| Visible | ***Human*** | a) Familiar (E1)<br>b) Non-familiar (E2) | E1/E2 sits behind the board, hands on the lap facing the mesh. |
| | ***Object*** | a) Film roll canister<br>b) Plastic animal<br>c) Police car | E2 sits behind the board, hands on her lap with the object placed in the middle of the board.<br>In the police car condition E2 holds the remote control and presses the horn button ten times. |
| | ***Food*** | a) Undesirable food<br>b) Dried fruit piece<br>c) 3 Raisins<br>d) Banana piece | E2 sits behind the board, hands on her lap with the food placed in the middle of the board. |
| | ***Non-Human*** | a) Red spot<br>b) Nothing | E2 places a red spot in the middle of the board and leaves.<br>Nothing is on the board and E2 is out of sight. |
| Movement | ***Human*** | a) Hand<br><br>b) Body | E2 sits behind the board and moves her right hand from the left side to the right side.<br>In the body condition E2 nods up and down while seating. |
| | ***Object*** | a) Film roll canister<br>b) Plastic animal<br>c) Police car | E2 sits behind the board and moves the object from the left side to right side and back on the board.<br>In the police car condition E2 lets the car drive to the other side of the board, left to right, two times. |
| | ***Food*** | a) Undesirable food<br>b) Dried fruit piece<br>c) 3 Raisins<br>d) Banana piece | E2 sits behind the board and moves the food from the left side to right side and back on the board. |
| Touch | ***Human*** | Hand | E2 sits behind the board and puts her right fist on the board. |
| | ***Object*** | a) Film roll canister<br>b) Plastic animal<br>c) Police car<br>d) Box | E2 sits behind the board, hands on her lap with the object placed on the board within reach of the individual. |
| | ***Food*** | a) Undesirable food<br>b) Dried fruit piece<br>c) 3 Raisins<br>d) Banana piece | E2 sits behind the board, hands on her lap with the food placed on the board within reach of the individual. |

### 2.1 Personality: Data analyses and detailed results

Results for the correlations between the three personality measures (latency, proximity and duration) and the performance in the two cognitive domains of the PCTB are reported in the main article. A multivariate analysis of variance of the three personality measures revealed no differences between ring-tailed and ruffed lemurs (Wilk's $\Lambda$=0.941, $F(3,34)$=0.71, p=0.550) and neither sex (Wilk's $\Lambda$=0.828, $F(3,34)$=2.35, p=0.090) nor the interaction between species and sex (species:sex; Wilk's $\Lambda$=0.955, $F(3,34)$=0.54, p=0.660) had an influence on individuals' performance. Univariate analyses (ANOVAs or Kruskal-Wallis-Tests) of each measure confirmed these insignificant differences between both species in all three personality measures.

### 2.2 Inhibitory Control Test

Testing inhibitory control of individuals (the ability to control one's impulses) might help to explain potential species differences in the physical or social domain of the tasks (Herrmann et al. 2007; Schmitt et al. 2012). It has been shown that inhibitory control can constrain apes in solving tasks in the physical (e.g. chimpanzees, Boysen and Berntson 1995) and the social domain (e.g. chimpanzees, Melis et al. 2006; Stevens and Hauser 2004). The inhibitory control test of this study consisted of six additional trials of the spatial memory task of the *PCTB* (experiment 1.1.1), assessing whether the individuals would skip the middle one out of three cups. Therefore, while the individual was watching, rewards were placed under the two outer cups and the middle cup was left empty. The individual could then choose one of the cups, and if it chose one of the baited cups correctly, it could choose a second time. No second choice was allowed if the individual chose the middle cup first. To correctly perform this task, individuals had to inhibit their tendency to choose the empty middle cup, which was positioned closest to them. Hence, a response was only scored as correct when the individual consecutively chose the two outer cups and skipped the middle cup. We found no differences in performance between the three species (Kruskal-Wallis test: $\chi^2$=2.34, p=0.31) and there were no correlations for inhibitory control and performance in any species, neither in the social nor the physical domain (Table S3).

**Table S3** Spearman rank correlation between inhibitory control and performance in the physical and social domain.

| species | domain | n | rho | p-value |
|---|---|---|---|---|
| Ruffed lemurs | physical | 13 | -0.13 | 0.662 |
| | social | 13 | -0.09 | 0.772 |
| Ring-tailed lemurs | physical | 27 | -0.08 | 0.691 |
| | social | 27 | -0.06 | 0.765 |
| Mouse lemurs | physical | 15 | -0.19 | 0.502 |
| | social | 15 | -0.12 | 0.677 |

### 3. Rank

The possible influence of the individuals' rank on performance was examined as well, except for the mouse lemurs that are housed solitarily. In all lemur groups, rank was inferred through additional focal animal observations.

### 4. Additional results

**Table S4** Comparisons of performance of the seven non-human primate species within the two domains. Presented are the results of *post hoc* multiple comparison analyses (Bonferroni); significant results are in boldface.

| | **physical domain** | **social domain** |
|---|---|---|
| Chimp - Orang | **<0.001** | 1 |
| Chimp - Baboon | 0.075 | 0.087 |
| Chimp - Macaque | **<0.001** | 0.842 |
| Chimp - Ruffed lemur | **<0.001** | 1 |
| Chimp - Ring-tailed lemur | **<0.001** | 1 |
| Chimp - Mouse lemur | **<0.001** | 1 |
| Orang - Baboon | 1 | **0.032** |
| Orang - Macaque | 0.700 | 0.269 |
| Orang - Ruffed lemur | 0.150 | 1 |
| Orang - Ring-tailed lemur | **<0.001** | 1 |
| Orang - Mouse lemur | **<0.001** | 1 |
| Baboon - Macaque | 1 | 1 |
| Baboon - Ruffed lemur | 1 | 0.291 |
| Baboon - Ring-tailed lemur | 0.070 | 0.207 |
| Baboon - Mouse lemur | 0.082 | 0.093 |
| Macaque - Ruffed lemur | 1 | 1 |
| Macaque - Ring-tailed lemur | 1 | 1 |
| Macaque - Mouse lemur | 1 | 0.927 |
| Ruffed lemur - Ring-tailed lemur | 1 | 1 |
| Ruffed lemur - Mouse lemur | 1 | 1 |
| Ring-tailed lemur - Mouse lemur | 1 | 1 |

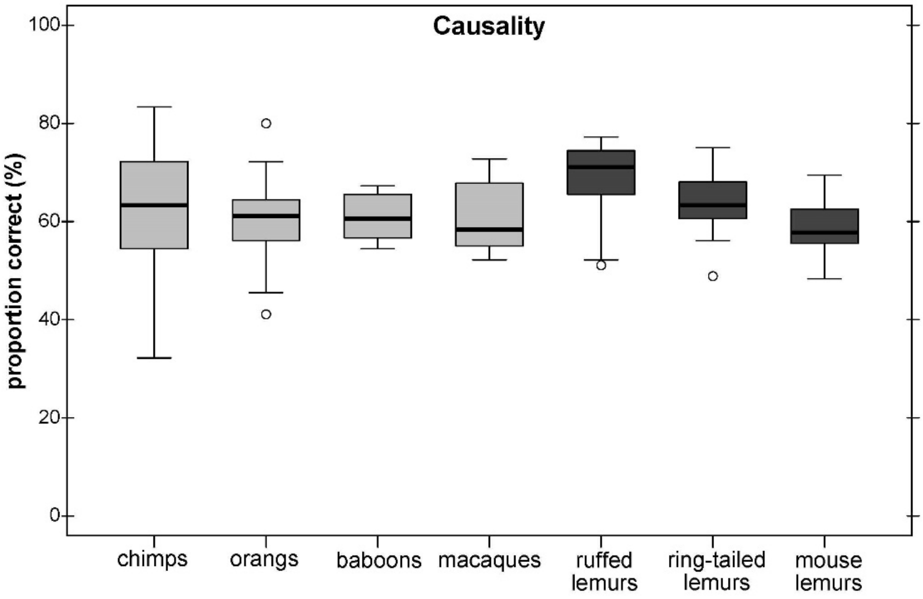


**Figure S2** Average performance of apes & monkeys (light grey) and lemurs (dark grey) in the scale causality excluding the tool use task. Represented are medians (black bars), interquartile ranges (boxes), upper and lower hinges (whiskers) and outliers (circles).
